# Angiogenic activity of cytochalasin B-induced membrane vesicles of human mesenchymal stem cells

**DOI:** 10.1101/646398

**Authors:** M.O. Gomzikova, M.N. Zhuravleva, V.V. Vorobev, I.I. Salafutdinov, A.V. Laikov, S.K. Kletukhina, E.V. Martynova, L.G. Tazetdinova, A.I. Ntekim, S.F. Khaiboullina, A.A. Rizvanov

## Abstract

**Background:** The cytochalasin B-induced membrane vesicles (CIMVs) are suggested to be used as a vehicle for the delivery of therapeutics. However, the angiogenic activity and therapeutic potential of human mesenchymal stem cells (MSCs) derived CIMVs (CIMVs-MSCs) remains unknown.

**Objectives:** The objectives of this study were to analyzed the morphology, size distribution, molecular composition and angiogenic properties of CIMVs-MSCs.

**Methods:** The morphology of CIMVs-MSC was analyzed by scanning electron microscopy. The proteomic analysis, multiplex analysis and immunostaining were used to characterize the molecular composition of the CIMVs-MSCs. The transfer of surface proteins from a donor to a recipient cell mediated by CIMVs-MSCs was demonstrated using immunostaining and confocal microscopy. The angiogenic potential of CIMVs-MSCs was evaluated using in vivo approach of subcutaneous implantation of CIMVs-MSCs in mixture with Matrigel matrix.

**Results:** Human CIMVs-MSCs retain parental MSCs content such as growth factors, cytokines, chemokines: EGF, FGF-2, Eotaxin, TGF-α, G-CSF, Flt-3L, GM-CSF, Fractalkine, IFNα2, IFN-γ, GRO, IL-10, MCP-3, IL-12p40, MDC, IL-12p70, IL-15, sCD40L, IL-17A, IL-1RA, IL-1a, IL-9, IL-1b, IL-2, IL-4, IL-5, IL-6, IL-7, IL-8, IP-10, MCP-1, MIP_1a, MIP-1b, TNF-α, TNF-β, VEGF. CIMVs-MSCs also have the expression of surface receptors similar to those in parental human MSCs (CD90^+^, CD29^+^, CD44^+^, CD73^+^). Additionally, CIMVs-MSCs could transfer membrane receptors to the surfaces of target cells *in vitro*. Finally, CIMVs-MSCs can induce angiogenesis *in vivo* after subcutaneous injection into adult rats.

**Conclusions:** Human CIMVs-MSCs have similar content, immunophenotype and angiogenic activity to those of the parental MSCs. Therefore, we believe that human CIMVs-MSCs could be used for cell free therapy of degenerative diseases.

## 1. Introduction

Broad differentiation potential of the stem cells (SCs) made them attractive tool for the regenerative medicine [1]. However, high risk of undesirable effects, including SCs malignant transformation [2, 3] and maldifferentiation [4, 5] limited their therapeutic applictaion. Therefore, the cell-free therapy concept was developed, where the SCs derived extracellular vesicles (EVs) are used instead [6].

EVs contain biologically active molecules of parental cells and mediate intercellular communication [7]. It was shown that the EVs can be used as a delivery vehicle as they have multiple advantages: 1) microvesicles retain the surface proteins of parental cells, 2) hydrophobic molecules (bioactive lipids and membrane proteins) could be transfered by EVs, 3) the EVs cytoplasmic membrane protects their content from degradation [6]. The regenerative potential of mesenchymal stem cells (MSCs) derived EVs was demonstrated using the rat model of myocardial infarction[8], where EVs restored the blood flow and reduced the size of the tissue damage [8]. MSC-EVs have also been reported to reduce the hepatic injury by improving the hepatocytes viability, suppressing oxidative injury and modulating the inflammatory response *in vivo* [9]. MSCs derived EVs were also successfully applied to treat transient global ischemia in mice, where the neuroprotective effects of MSC-EVs was demonstrated [10].

The potential of Cytochalasin B-induced membrane vesicles (CIMVs) as a vector for drug delivery has been demonstrated [11]. We have previously shown that CIMVs have size similar to that of the natural EVs [12]. CIMVs also retain the biological activity of parental cells and are able to stimulate capillary tube formation *in vitro* and vasculogenesis *in vivo* [12]. These properties of CIMVs led to our interest in the evaluation of the biological activity of MSCs derived CIMVs (CIMVs-MSCs) as this might provide a potential tool for cell-free therapy.

The objectives of this study were to determine the morphology, molecular content and angiogenic activity of human CIMVs-MSCs as well as describe the membrane receptor transferring and biological activity of human CIMVs-MSCs.

## 2. Materials and methods

### 2.1 MSCs isolation and characterization

Human sample collection was approved by the local Ethical Committee of Kazan (Volga region) Federal University based on article 20 of the Federal Legislation on “Health Protection of Citizens of the Russian Federation” № 323-FL, 21.11.2011. Signed informed consent was obtained from each donor. To obtain cell suspension the adipose tissue was cut into small pieces and treated with 0.2% collagenase II (Dia-M, Russia) in a shaker-incubator at 37 °C, 120 rpm for one hour. Cell suspension was pelleted (400g for 5 min), washed once in PBS (PanEco, Russia) and re-suspended in DMEM (PanEco, Russia) supplemented with 10% fetal bovine serum (Gibco, UK) and 2mM L-glutamine (PanEco, Russia). MSCs were maintained at 37°C, 5% CO_2_ with culture medium replaced every three days.

MSCs were differentiated into three directions: adipogenic, chondrogenic and osteogenic. Adipogenic, chondrogenic and osteogenic differentiation was confirmed by detection of lipid droplets (Oil Red dye staining), glycosaminoglycans and mucins (1% alcian blue staining) and calcium deposits (5% AgNO_3_ staining), respectively [13].

The immune phenotype of isolated cells was analyzed by staining with monoclonal antibodies CD90-PE/Cy5 (328112; BioLegend, USA), CD90- Brilliant Violet 421 (328122; BioLegend, USA); CD44-APC/Cy7 (103028; BioLegend, USA), CD29-APC (2115040; Sony, USA), CD73- APC (51-9007649; BD bioscience, USA), CD73-PerCP-Cy5.5 (344014; BioLegend, USA), STRO-1- APC/Cy7 (340104; BioLegend, USA), CD45-FITC (304006; BioLegend, USA). Expression of CD markers were analyzed by flow cytometry using BD FACS Aria III (BD bioscience, USA).

### 2.2 CIMVs production

CIMVs were prepared as described previously [12]. Briefly, MSCs were washed twice with PBS, and maintained in DMEM supplemented with 10 μg/ml of Cytochalasin B (Sigma-Aldrich, USA) for 30 min (37°C, 5% CO_2_). Cell suspension was vortexed vigorously for 30 sec and pelleted (100 g for 10 min). The supernatant was collected and subject to two subsequent centrifugation steps (100g for 20 min and 2000g for 25 min). The pellet from last step, containing CIMVs-MSC, was washed once in PBS (2000g for 25 min).

### 2.3 Characterization of the CIMVs

#### Scanning electron microscopy (SEM)

CIMVs were fixed (10% formalin for 15 min) and dehydrated using graded alcohol series and dried at 37°C. Prior to imaging, samples were coated with gold/palladium in a Quorum T150ES sputter coater (Quorum Technologies Ltd, United Kingdom). Slides were analyzed using Merlin field emission scanning electron microscope (CarlZeiss, Germany). For the size analysis, three independent batches of CIMVs were produced and used to generate at least six electron microscope images for each batch. Data collected was used to determine the CIMVs size.

### Proteome analysis

CIMVs-MSCs and MSCs were lysed in RIPA buffer (150 mM NaCl, 1% NP-40, 0.5% sodium deoxycholate, 0.1% SDS, 25 mM Tris (pH 7.4)) and separated using gel polyacrylamide gel electrophoresis [14]. Gels were fixed overnight (20% ethanol and 10% acetic acid), strips were cut (1.5×1.5 mm), dehydrated using 100% acetonitrile for 20 min and dried at the room temperature. Gel fragments were rehydrated (200 mM ammonium bicarbonate, 100% acetonitrile, dH2O), placed in sequencing grade-modified trypsin (Promega, USA) and incubated overnight at 37 °C. The cleaved peptides were extracted from the gel pieces using extraction buffer (0.5% trifluoroacetic acid (TFA)) and incubated in an ultrasonic bath for 10 minutes, followed by adding 100% acetonitrile and 0.5% TFA. The mixture of peptides was dried at 45°C under vacuum using Concentrator plus Complete System (Eppendorf, USA). Samples were desalted using the Acclaim PepMap 100 Columns (160321, Thermo Scientific, USA) (C18, 3 μm, 100Å) for 5 min at a 5 μl/min flow rate.

Liquid chromatography mass spectrometry analysis (LC-MS / MS) of peptide extracts was done using 3000 Nano LC nanochromatographic system (Thermo Scientific, USA) and Maxis Impact mass spectrometer with an electrospray ionization source Captive Spray (Bruker, USA). Peptides were separated by reverse phase chromatography using an Acclaim PepMap 100 NanoViper column (C18, 2μm, 100Å, 75μm × 15cm) (Thermo Scientific, USA). The tryptic peptides were eluted in a linear gradient with a mixture of solution A (4.5% acetonetrile in diH 2 O with 0.5% formic acid) and increasing percentage (from 5 to 35%) of solution B (94.5% acetonetrile with 0.5% formic acid). Elution was done for 60 minutes at 40°C and a 300 nl/min flow rate, 1600 V of the Captive Spray source, capillary temperature 150°C, 3.0 l/min dry gas flow. The positive polarity and total spectrum measurements, data dependent acquisition (DDA) were set on Maxis Impact mass spectrometer (Bruker, USA). Peptides mass spectrum was compared to the theoretical peptide masses of all human proteins using the SWISS-PROT and NCBI databases.

### Multiplex analysis

Multiplex analysis based on the xMAP Luminex technology was performed with the use of MILLIPLEX MAP Human Cytokine/Chemokine Magnetic Bead Panel - Premixed 38 Plex - Immunology Multiplex Assay (sCD40L, EGF, Eotaxin/CCL11, FGF-2, Flt-3 ligand, Fractalkine, G-CSF, GM-CSF, GRO, IFN-α2, IFN-γ, IL-1α, IL-1β, IL-1ra, IL-2, IL-3, IL-4, IL-5, IL-6, IL-7, IL-8, IL-9, IL-10, IL-12 (p40), IL-12 (p70), IL-13, IL-15, IL-17A, IP-10, MCP-1, MCP-3, MDC/CCL22, MIP-1α, MIP-1β, TGF-α, TNF-α, TNF-β, VEGF) (Merckmillipore, USA), in accordance with the manufacturer’s instructions. Briefly, samples were incubated with fluorescent beads for 1 hour, washed and incubated with phycoerythrin-streptavidin for 10 min (Merckmillipore, USA). The analysis was done using a Luminex 200 analyzer (Merckmillipore, USA). The CIMVs-MSCs and MSCs lysates in IP buffer (50 mMTris-Cl, 150 mMNaCl, 1% Nonidet-P40) were used for multiplex analysis. Equal protein load was used the analysis.

#### Flow cytometry analysis

The immune phenotype of CIMVs-MSCs was analyzed by immunostaining with monoclonal antibodies: CD90-PE-Cy5 (328112; BioLegend, USA), CD29- APC (2115040; Sony, USA), CD44-APC/Cy7 (103028; BioLegend, USA), CD73-PerCP/Cy5.5 (344014; BioLegend, USA). CIMVs were analyzed by flow cytometry (BD FACS Aria III. BD Bioscience, USA), the 405 nm laser was used for better resolution of CIMVs-MSC.

#### Cytoplasmic membrane staining

CIMVs were stained with lipophilic dye DiD (V22889; Life Technologies, USA) according to the manufacturer’s instructions. Briefly, CIMVs (300 μg/ml) were incubated in 5μM of DiD dye for 15 min (37°C, 5% CO_2_) and washed twice with complete medium (DMEM with 10% FBS, 2mM L-glutamine) before use.

### 2.4 Animals

Adult rats (Rattus norvegicus) (Pushchino, Russia) were used. All experiments were carried out in compliance with the procedure protocols approved by KFU local ethics committee (protocol #5, date 27.05.2014) according to the rules adopted by KFU and Russian Federation Laws. All experiments were repeated three times. Rats were euthanized using CO_2_ in compliance with the procedure protocols approved by KFU local ethics committee (protocol #5, date 27.05.2014).

### 2.5 Angiogenic activity test in vivo

Rats were injected with MSCs (1×10^6^) or CIMVs (50 μg) in 400 μl Matrigel Basement Membrane Matrix (10 mg/ml) (356231, BectonDickinson, USA) subcutaneously. Animals were injected with MSCs and CIMVs-MSCs unstained or prestained with DiO membrane dye (V-22886, LifeTechnoligies, USA). Control animals were injected with the Matrigel matrix (10 mg/ml). Eight days later, fragments of Matrigel matrix were collected, fixed with 10% buffered formalin solution (BioVitrum, Russia), dehydrated in an ethanol gradient and embedded into Histomix paraffin (BioVitrum, Russia). Paraffin sections (6 μm thick) were cut using HM 355S microtome (ThermoScientific, USA), dewaxed with Roticlear (CarlRoth, USA) and stained with hematoxylin-eosin (BioVitrum, Russia). Then the sections were dehydrated and embedded in a Canadian balsam (PanReac AppliChem, Spain). Slides were examined using AxioOberver.Z1 (CarlZeiss, Germany) fluorescent microscope. Five fragments of Matrigel matrix and at least three microscope images were examined for each batch.

Fragments of Matrigel matrix containing MSCs or CIMVs prestained with DiO dye (V-22886, LifeTechnoligies, USA) were stored in liquid nitrogen. Matrigel matrix slides (6 μm thick) were made using HM560 Cryo-Star microtome (ThermoScientific, USA). Nucleus was stained using 5 μg/ml DAPI (D1306, Invitrogen, USA).

### 2.6 Statistical analysis

Statistical analysis was done using Wilcoxon signed-rank test (R-Studio) with significance level p<0.05. Illustrations were built with “ggplot2” package (v3.1.0, 2018).

## 3. RESULTS

### 3.1 Isolation and characterization of human adipose-derived MSCs

Primary MSCs were isolated from human subcutaneous adipose tissue. MSCs phenotype was confirmed using antibodies against CD90, CD73, CD44, CD105 and CD45 (Fig.1 A). Next, differentiation potential (chondrogenic, adipogenic and osteogenic) of isolated MSCs were analyzed (Fig.1 B).

**Figure 1.**
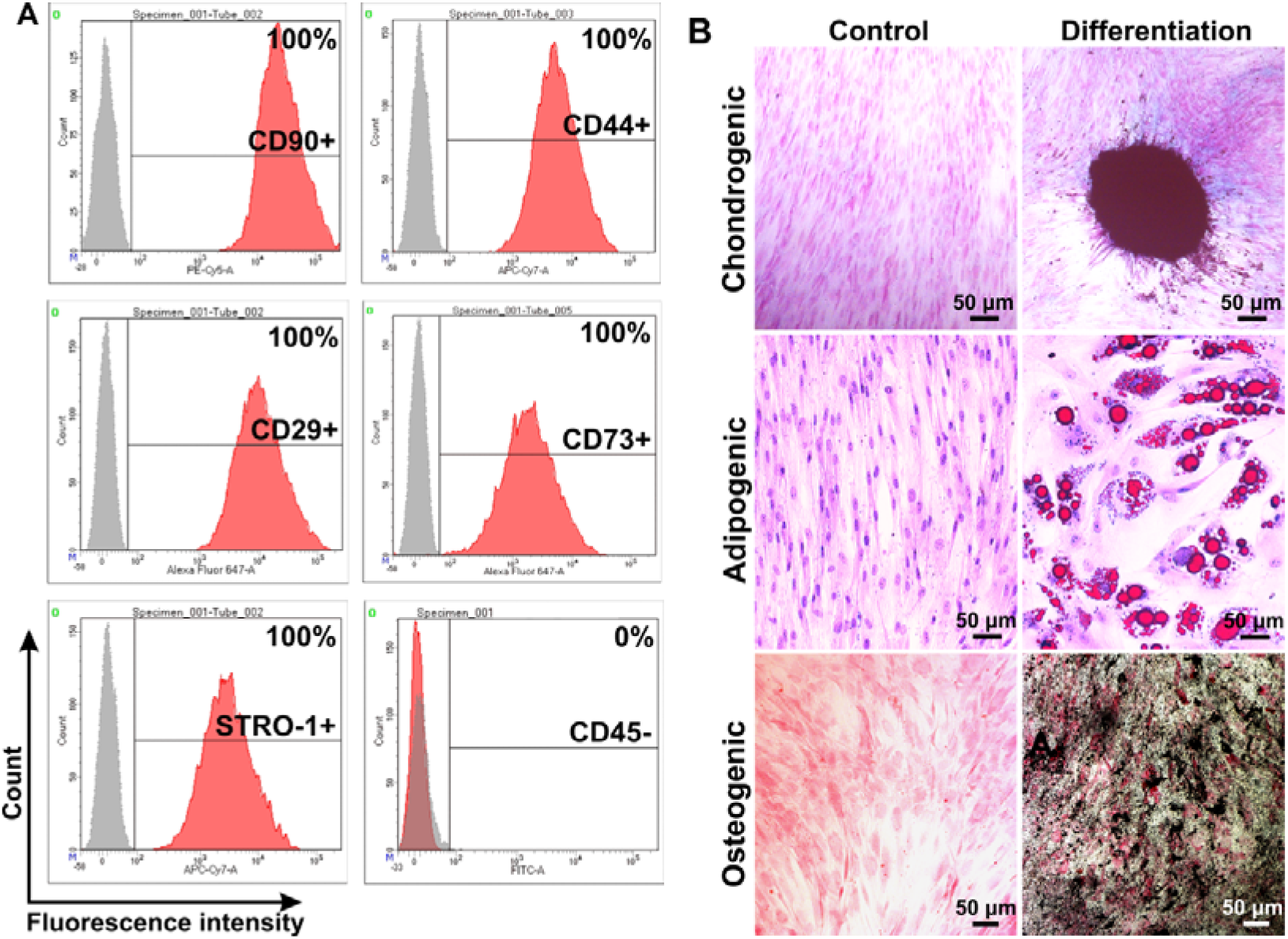
Phenotype analysis of human MSCs isolated from subcutaneous adipose tissue. Flow cytometry data (A). Histograms were generated using FACSDiva7 software (BDBioscience, USA). Grey– negative control; Red –cells labeled with antibodies. Analysis of MSCs differentiation into: chondrogenic, adipogenic and osteogenic progenitors (B). Images were captured using ZEISS Axio Observer Z1 microscope (CarlZeiss, Germany).

Cells phenotype were determined as CD90^+^, CD44^+^, CD29^+^, CD73^+^, STRO-1^+^ and CD45^−^ (Fig.1 A), which are characteristics of the MSCs [15]. Isolated cells could be differentiated into chondrogenic, adipogenic and osteogenic progenitors confirming the multipotency of isolated MSCs (Fig.1 B).

### 3.2 Characterization of human CIMVs-MSCs

CIMVs were successfully generated from primary human adipose MSCs (Fig.2). The morphology and size of the human CIMVs-MSCs were analyzed using scanning electron microscopy (SEM). CIMVs-MSCs had spherical structures and sizes ranging from 100 to 2600 nm with the majority (89.36%) having sizes between 100-1200 nm (Fig.2 B).

**Figure 2.**
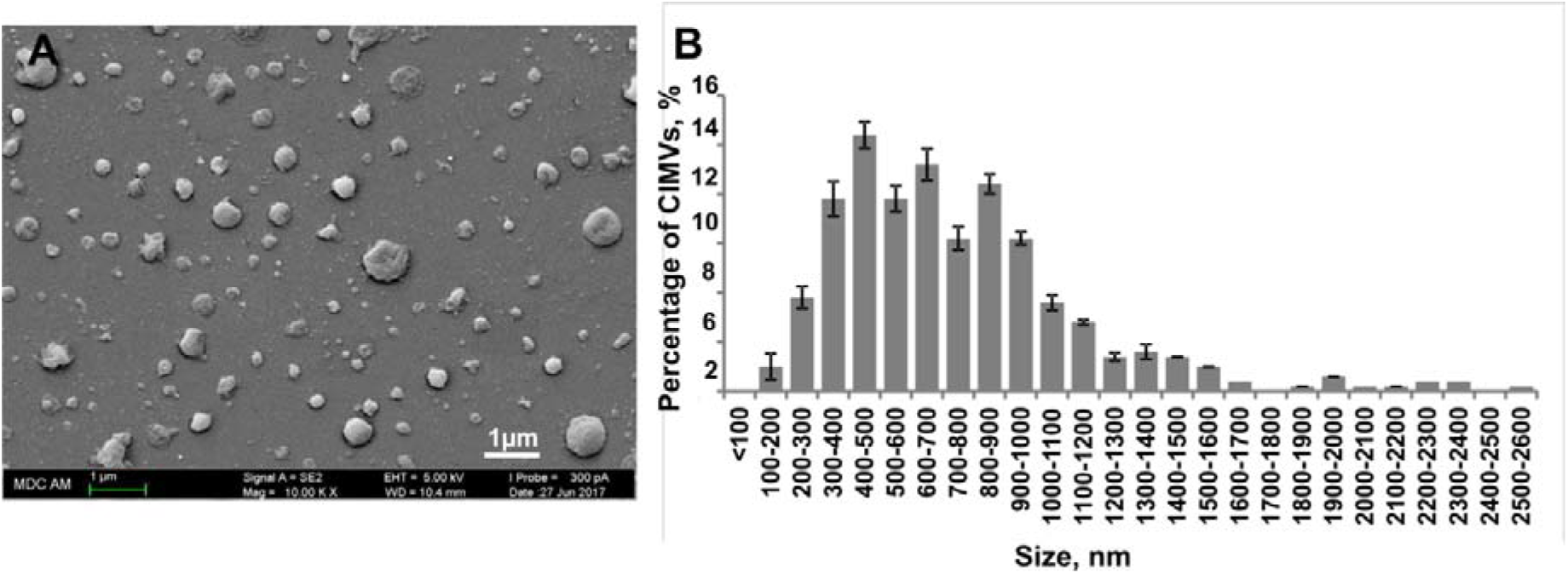
Analysis of the morphology and the size distribution of human CIMVs-MSCs. Human CIMVs-MSCs were characterized using scanning electron microscopy (A). At least six electron microscope images were analyzed from three independent experiments to determine the size of human CIMVs-MSCs (B).

Molecular composition of human CIMVs-MSCs was examined using proteome and xMap Luminex multiplex analysis (Fig. 3, 4). Proteome analysis identified 373 proteins in human MSCs and 362 proteins in CIMVs-MSCs lysates. Interestingly, the majority (252 molecules) of proteins were similar between MSCs and CIMVs-MSCs while 121 (32.4%) and 110 (30.4%) proteins were unique in MSC and CIMVs-MSCs, respectively (Fig. 3 A).

**Figure 3.**
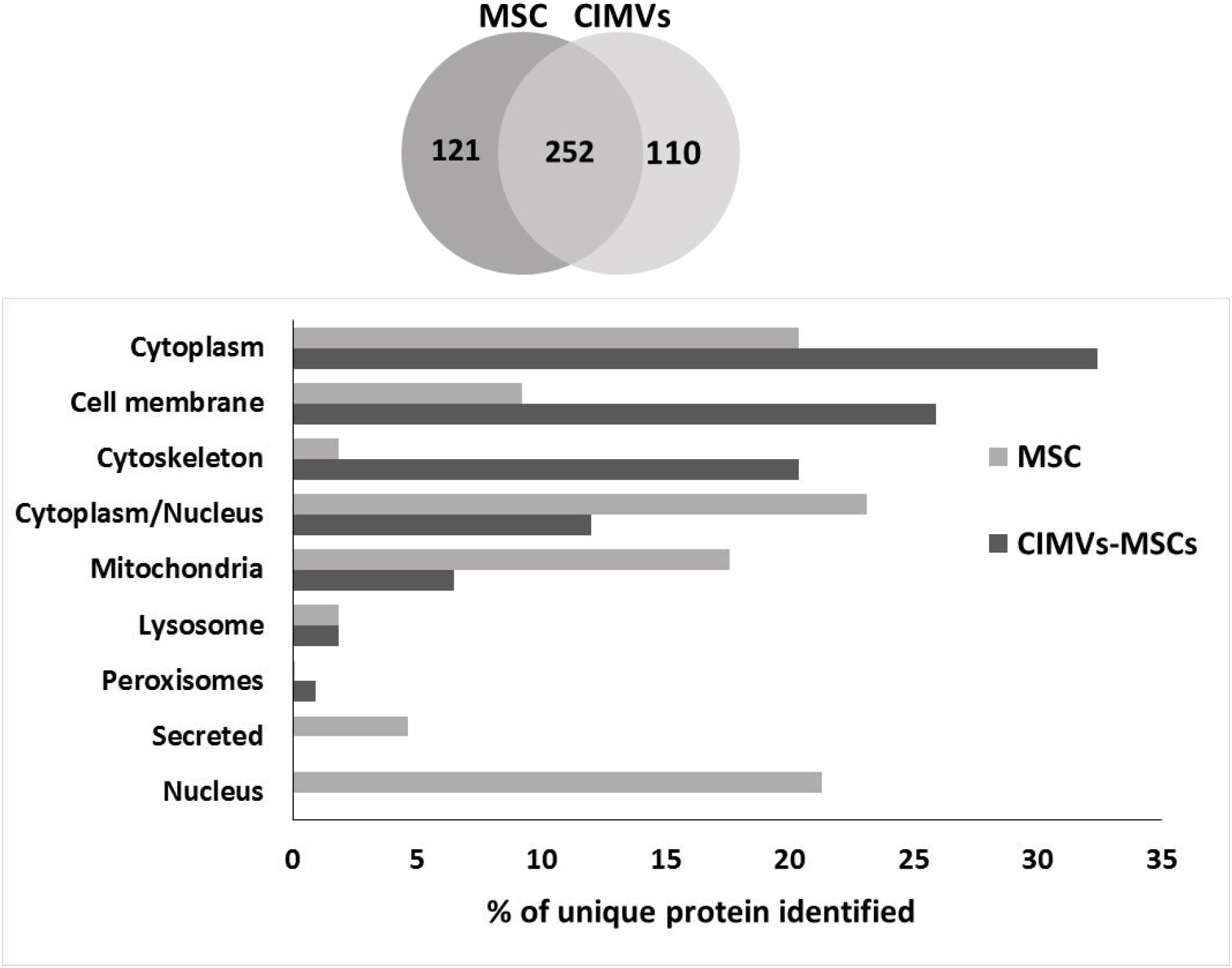
Proteome analysis of human MSCs and CIMVs-MSCs. Venn diagram of identified proteins MSCs and CIMVs-MSCs (A). Distribution of the identified proteins in organelles, % of unique identified proteins (B).

**Figure 4.**
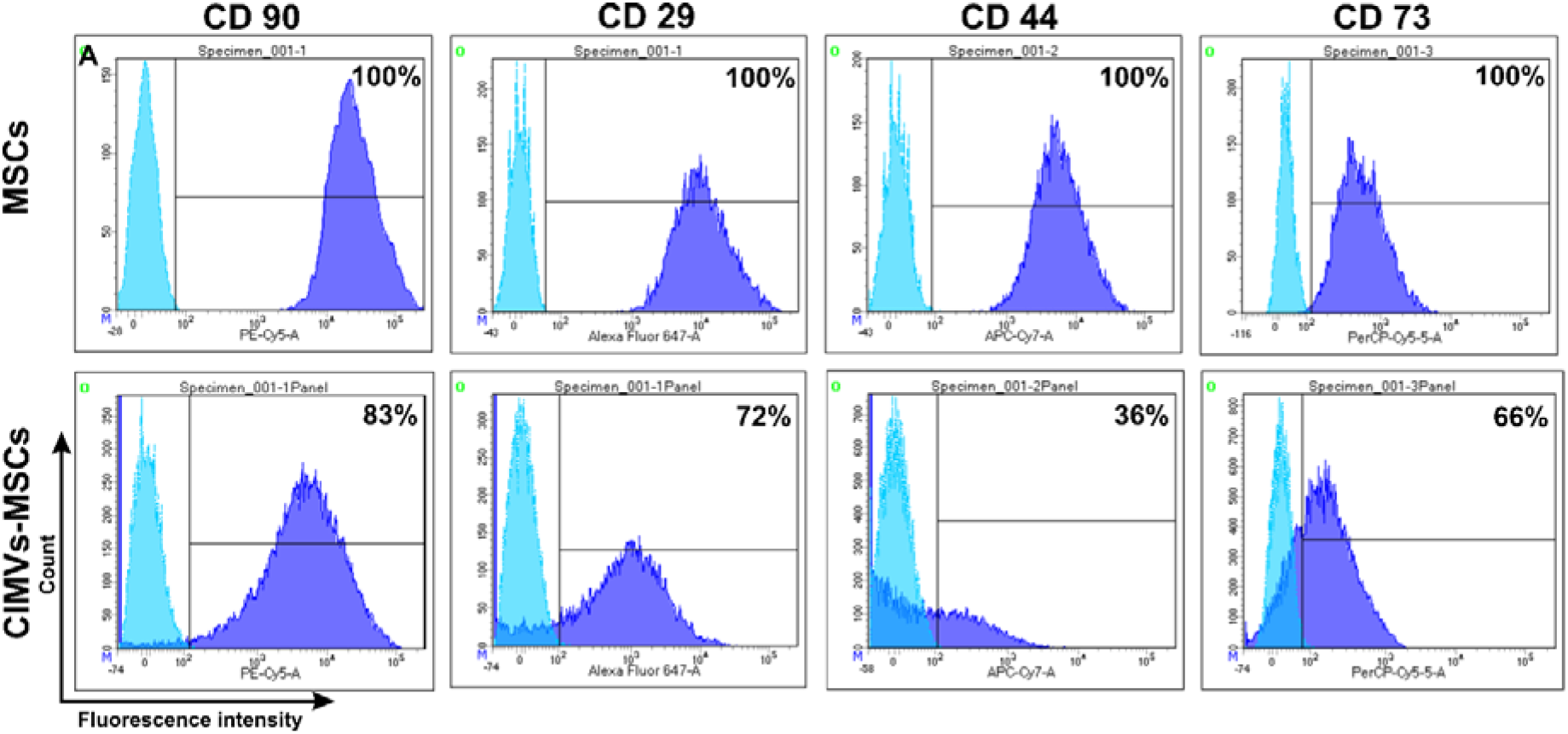
Immune phenotype of human MSCs and CIMVs-MSCs. MSCs and CIMVs-MSCs were stained with anti-CD90, anti-CD29, anti-CD44, anti-CD73 monoclonal antibodies and analyzed using flow cytometer BD FACS Aria III (BD Bioscience, USA). Histograms were generated using FACSDiva7 software (BD Bioscience, USA). Blue – isotype control; Dark blue – MSCs or CIMVs-MSCs labeled with antibodies.

The unique proteins in human MSCs included nuclear (21.3%), secreted (4.6%), lysosomal (1.9%), mitochondrial (17.6%), cytoplasmic/nuclear (23.1%), cytoskeleton (1.8%), cell membrane (9.3%) and cytoplasm (20.4%) associated (Fig. 3 B). Peroxisome proteins were below the proteomics detection range in MSCs.

The unique proteins in CIMVs-MSCs included proteins associated with peroxisome (0.9%), lysosome (1.8%), mitochondria (6.5%), cytoplasm/nucleus (12%), cytoskeleton (20.4%), cell membrane (26%) and cytoplasm (32.4%) (Fig. 3 B). Nuclear and secreted proteins were below the proteomic detection range in CIMVs-MSCs.

A multiplex approach was used to characterize the molecular content of CIMVs-MSCs. Multiple cytokines were similar between CIMVs-MSCs and parental MSCs. These included growth factors, cytokines and chemokines (EGF, FGF-2, Eotaxin, TGF-α, G-CSF, Flt-3L, GM-CSF, Fractalkine, IFNα2, IFN-γ, GRO, IL-10, MCP-3, IL-12p40, MDC, IL-12p70, IL-15, sCD40L, IL-17A, IL-1RA, IL-1a, IL-9, IL-1b, IL-2, IL-4, IL-5, IL-6, IL-7, IL-8, IP-10, MCP-1, MIP_1a, MIP-1b, TNF-α, TNF-β and VEGF) (Table 1). Levels of IL-3 and IL-13 were below the detection range in MSCs and CIMVs-MSCs. Interestingly, levels of TGF-β, CCL7, sCD40L, IL-1b and TNF-β were higher in MSCs as compared to CIMVs-MSCs.

**Table 1.**
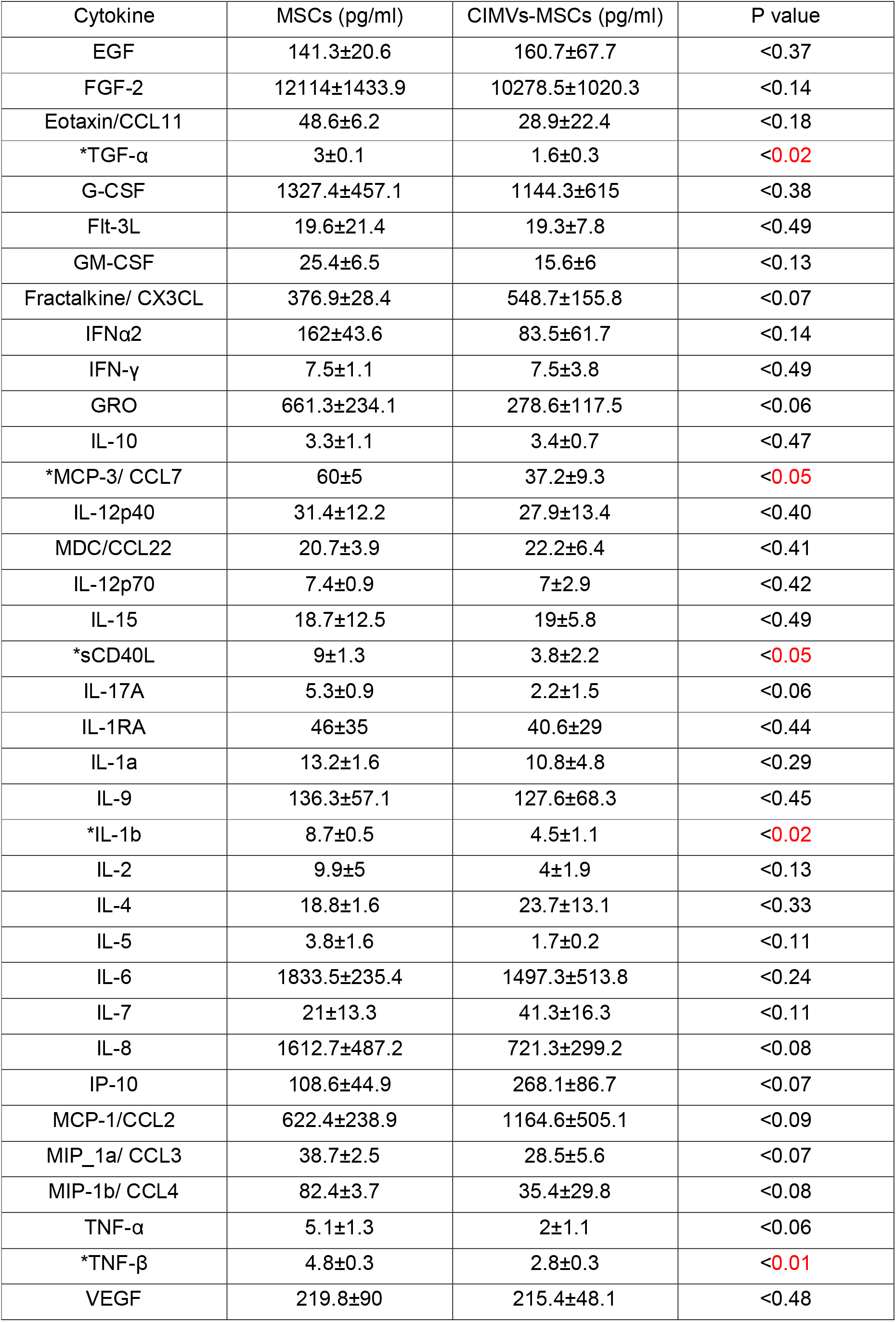
Cytokine analysis of MSCs and CIMVs-MSCs content.

| Cytokine | MSCs (pg/ml) | CIMVs-MSCs (pg/ml) | P value |
| --- | --- | --- | --- |
| EGF | 141.3±20.6 | 160.7±67.7 | <0.37 |
| FGF-2 | 12114±1433.9 | 10278.5±1020.3 | <0.14 |
| Eotaxin/CCL11 | 48.6±6.2 | 28.9±22.4 | <0.18 |
| *TGF-α | 3±0.1 | 1.6±0.3 | <0.02 |
| G-CSF | 1327.4±457.1 | 1144.3±615 | <0.38 |
| Flt-3L | 19.6±21.4 | 19.3±7.8 | <0.49 |
| GM-CSF | 25.4±6.5 | 15.6±6 | <0.13 |
| Fractalkine/ CX3CL | 376.9±28.4 | 548.7±155.8 | <0.07 |
| IFNα2 | 162±43.6 | 83.5±61.7 | <0.14 |
| IFN-γ | 7.5±1.1 | 7.5±3.8 | <0.49 |
| GRO | 661.3±234.1 | 278.6±117.5 | <0.06 |
| IL-10 | 3.3±1.1 | 3.4±0.7 | <0.47 |
| *MCP-3/ CCL7 | 60±5 | 37.2±9.3 | <0.05 |
| IL-12p40 | 31.4±12.2 | 27.9±13.4 | <0.40 |
| MDC/CCL22 | 20.7±3.9 | 22.2±6.4 | <0.41 |
| IL-12p70 | 7.4±0.9 | 7±2.9 | <0.42 |
| IL-15 | 18.7±12.5 | 19±5.8 | <0.49 |
| *sCD40L | 9±1.3 | 3.8±2.2 | <0.05 |
| IL-17A | 5.3±0.9 | 2.2±1.5 | <0.06 |
| IL-1RA | 46±35 | 40.6±29 | <0.44 |
| IL-1a | 13.2±1.6 | 10.8±4.8 | <0.29 |
| IL-9 | 136.3±57.1 | 127.6±68.3 | <0.45 |
| *IL-1b | 8.7±0.5 | 4.5±1.1 | <0.02 |
| IL-2 | 9.9±5 | 4±1.9 | <0.13 |
| IL-4 | 18.8±1.6 | 23.7±13.1 | <0.33 |
| IL-5 | 3.8±1.6 | 1.7±0.2 | <0.11 |
| IL-6 | 1833.5±235.4 | 1497.3±513.8 | <0.24 |
| IL-7 | 21±13.3 | 41.3±16.3 | <0.11 |
| IL-8 | 1612.7±487.2 | 721.3±299.2 | <0.08 |
| IP-10 | 108.6±44.9 | 268.1±86.7 | <0.07 |
| MCP-1/CCL2 | 622.4±238.9 | 1164.6±505.1 | <0.09 |
| MIP_1a/ CCL3 | 38.7±2.5 | 28.5±5.6 | <0.07 |
| MIP-1b/ CCL4 | 82.4±3.7 | 35.4±29.8 | <0.08 |
| TNF-α | 5.1±1.3 | 2±1.1 | <0.06 |
| *TNF-β | 4.8±0.3 | 2.8±0.3 | <0.01 |
| VEGF | 219.8±90 | 215.4±48.1 | <0.48 |

### 3.3 Immunophenotype of human CIMVs-MSCs

MSCs surface receptors play role in cell to cell contact, immunomodulation and activation of signaling in target cells [16]. Therefore, we sought to determine if CIMVs retain the surface receptors of MSCs. All parental MSCs (100%) expressed CD90, CD29, CD44 and CD73 (Fig.4) characteristic for the MSCs [15]. CIMVs-MSCs were positive for CD90, CD29, CD44 and CD73 (83%, 72%, 36% and 66%, respectively) (Fig.4).

### 3.4 Transfer of cell surface receptors to the recipient cell membrane by CIMVs-MSCs

Microvesicles can transfer soluble factors as well as surface receptors by the fusion of cytoplasmic membranes [17, 18]. Therefore, we sought to determine whether CIMVs-MSCs could transfer the surface receptors to the recipient HEK293FT cells. HEK293FT cells were pre-stained with DiO (Invitrogen, USA) and cultured for 24 hours with DiD labeled CIMVs-MSCs (10μг/мл) (Invitrogen, USA). Expression of CD90 was selected to demonstrate receptor transfer, as it is specific for CIMVs-MSCs and absent on HEK293FT cells. Expression of CD90 was analyzed using laser scanning confocal microscope Zeiss LSM 780 (Carl Zeiss, Germany) and flow cytometry BD FACS Aria III (BD Bioscience, USA). We found that CIMVs-MSCs and HEK293FT membranes became fused and CD90 surface receptor was transferred to HEK293FT (Fig.5 D-G). We determined that 99.14% of HEK293FT recipient cells acquired CD90+ immunophenotype (Fig.5 H, I).

**Figure 5.**
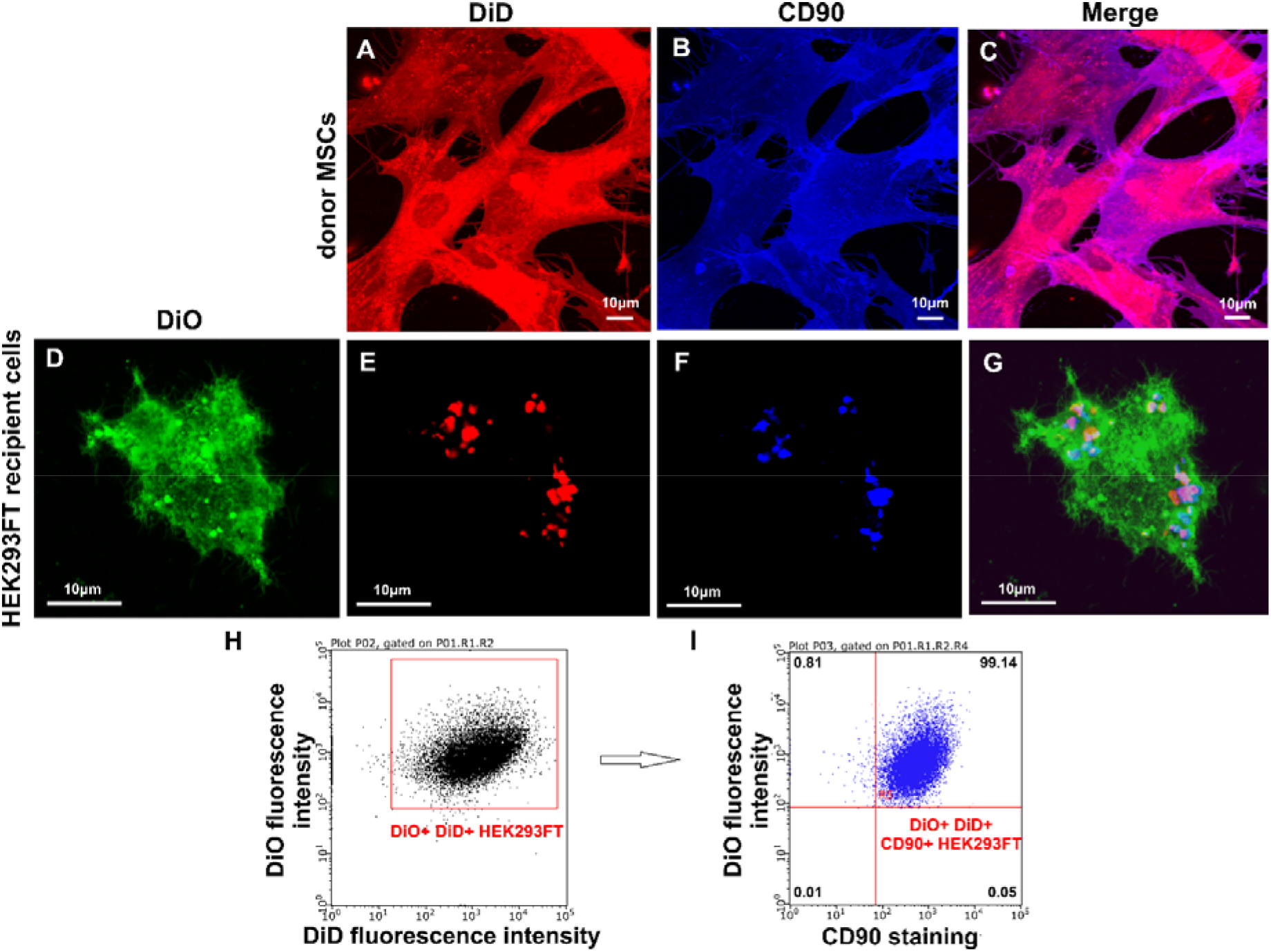
Analysis of CD90 transfer by CIMVs-MSCs to recipient HEK293FT cells. A-G - laser scanning confocal microscopy using Zeiss LSM 780 (Carl Zeiss, Germany), H-I - flow cytometry using BD FACS Aria III (BD Bioscience, USA).

### 3.5 CIMVs-MSCs stimulated angiogenesis *in vivo*

Since the CIMVs-MSCs contain multiple growth factors (EGF, FGF-2 and VEGF), we postulated that CIMVs-MSCs could have angiogenic activity. We used *in vivo* approach to demonstrate angiogenetic capacity of MSCs and CIMVs-MSCs. MSCs (1×10^6^ cells) and CIMVs-MSCs (50 μg) were stained with vital membrane dye DiO (Invitrogen, USA), mixed with Matrigel matrix (400 μl) and injected into rats subcutaneously. Eight days later, MSCs and CIMVs-MSCs containing Matrigel matrix plugs were collected from the subcutaneous space of the rats and fixed in 10% formalin (group 1) or frozen in liquid nitrogen (group 2). Formalin-fixed Matrigel matrix plugs were stained with hematoxylin-eosin kit (BioVitrum, Russia) (Fig.6 A-C). Frozen Matrigel matrix plugs were cut using HM560 Cryo-Star microtome (ThermoScientific, USA) and stained with DAPI (D1306, Invitrogen, USA) (Fig.6 D-F).

**Figure 6.**
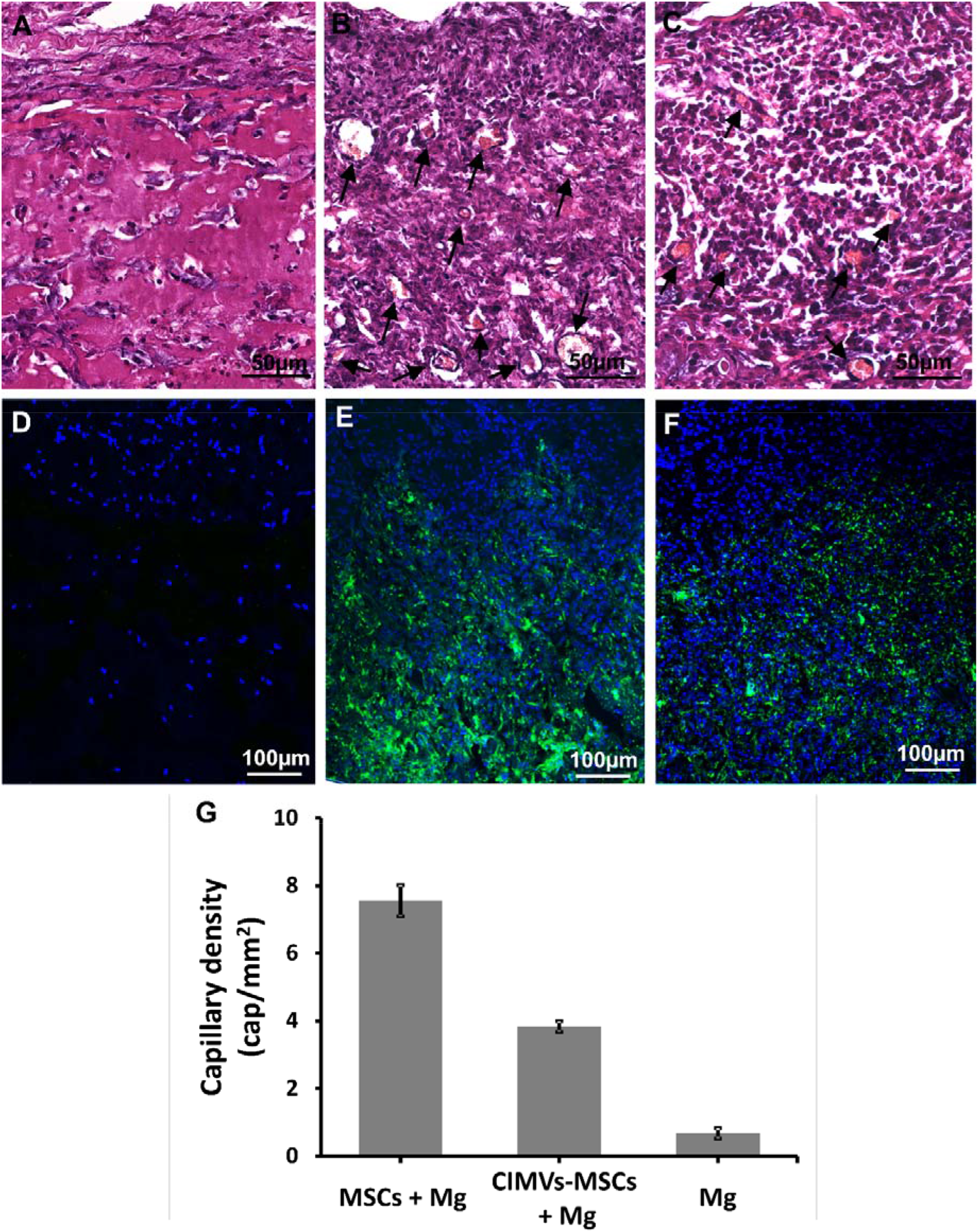
Analysis of angiogenic activity of MSCs and CIMVs-MSCs *in vivo*. Hematoxylin/eosin staining and fluorescence micrographs are shown after subcutaneous injections in rats (6 animals per experimental group) of MSCs (10^6^ cells) or CIMVs-MSCs (50 μg). A, D – negative control (s.c. injection of Matrigel matrix); B, E – s.c. injection of MSCs, C,F - s.c. injection of CIMVs-MSCs. Arrows mark the position of the sprouting blood capillaries. Counting the total number of vessels was carried out using the AxioVision 4.8 program (CarlZeiss). (G) Quantitation of the capillary density in Matrigel matrix plugs. The data represents mean ± SD. For statistical analysis, ten hematoxylin and eosin stained slides per animal were analyzed.

MSCs and CIMVs-MSCs were detected in the Matrigel matrix implants 8 days after s.c. injection (Fig 6 D-F). Also, newly developed blood capillaries were observed in Matrigel matrix containing MSCs and CIMVs-MSCs (Fig 6 A-C). We found that the number of the newly developed blood vessels in control Matrigel matrix (without MSCs or CIMVs) was 0.67 ± 0.15 cap/mm^2^ (Fig.6 A, D, G). In Matrigel matrix containing MSCs, the number of newly developed blood vessels was 11.3-fold higher (7.55 ± 0.46 cap/mm^2^, p<0.01) than that in control (Fig.6 B, E, G). Similar to MSCs, the number of the new capillaries in Matrigel matrix containing CIMVs-MSCs was increased (5.7-fold higher (3.84 ± 0.16 cap/mm^2^, p<0.01)) than that in control (Fig.6 C, F, G). We suggest that the human CIMVs-MSCs retain the angiogenic activity of the parental MSCs.

## 4. Discussion

MSCs derived EVs retain angiogenic and regenerative properties of the parental cells [19, 20]. The yield of the naturally produced EVs by MSCs is low, limiting their clinical application. Large scale EVs production was made using Cytochalasin B [21]. CIMVs could be produced in large quantities while retaining the size of natural EVs [12, 21]. However, our understanding of the biologic activity and therapeutic potential of MSCs derived CIMVs remains limited.

Here, for the first time, we have shown that the size of the majority of human CIMVs-MSCs ranges between 100 and 1200 nm (89.36%), which is similar to that of EVs. Also, the proteome content of human MSCs and CIMVs-MSCs appears to be similar. Analysis of the CIMVs-MSCs content revealed an increased proteins linked to cytoskeleton, peroxisomes, cell membrane and cytoplasm. In contrast, mitochondria and cytoplasm/nucleus proteins were decreased, while nucleus and secreted proteins were significantly depleted as compared to MSCs. We believe that the cytoskeleton proteins and membrane proteins enrichment of CIMVs-MSCs is due to the mechanism of their outward release from the cell surface. Similar data was demonstrated by Kim and colleagues [22]. We suggest that the enrichment of peroxisomes/cytoplasm proteins and depletion of mitochondria/cytoplasm/nucleus proteins could be due to the deep intracellular localization of these organelles.

MSCs activate cell proliferation, migration and angiogenesis in vivo by direct contact and paracrine mechanisms including secretion of growth factors, cytokines and chemokines [23]. Therefore, we sought to analyze the cytokine content and immune phenotype of human CIMVs-MSCs. We found that CIMVs-MSCs have the molecular content similar to that in parental MSCs (Table 1). Interestingly, CIMVs-MSCs had significantly lower level of TGF-α (<0,02), MCP-3/ CCL7 (<0,05), sCD40L (<0,05), IL-1b (<0,02) and TNF - β (<0,01). Levels of growth factors and interleukins in CIMVs-MSCs were similar to that in MSCs (Table 1). To our knowledge, this is the first time the cytokines content of CIMVs-MSCs have been characterized. There is limited data on cytokine content of MSCs available. Several cytokines were detected by Mussano F and colleagues. in MSCs culture medium. These included IL-2, IL-6, IL-8, IL10, IL-12, G-CSF, INF-γ, TNF-α, MCP-1 (CCL-2), IP-10, PDGF, bFGF and VEGF [24]. The authors reported that MSCs produced high levels of IL-6, IL-8, MCP-1 (CCL-2) and VEGF [24]. Schinkothe and colleagues reported that human MSCs produced high level of G-CSF, IL-12p40, IL-17, CCL2, CCL3 and CCL4 [25]. These data corroborate our results, where we have detected FGF2/ bFGF, G-CSF, IFN-γ, IL-10, IL-12p40, IL-17A, IL-2, IL-6, IL-8, IP-10, MCP-1/CCL2, MIP_1a/ CCL3, MIP-1b/ CCL4, TNF-α and VEGF in human MSCs and CIMVs-MSCs. The presence of several other cytokines in human MSCs and CIMVs-MSCs were demonstrated in our study, these include EGF, Eotaxin/CCL11, TGF-α, Flt-3L, GM-CSF, Fractalkine/ CX3CL, IFNα2, GRO, MCP-3/ CCL7, MDC/CCL22, IL-12p70, IL-15, sCD40L, IL-1RA, IL-1a, IL-9, IL-1b, IL-4, IL-5, IL-7 and TNF-β.

We found that CIMVs-MSCs have the surface receptors similar to that of the parental human MSCs: CD90^+^ (83%), CD29^+^ (72%), CD44^+^ (36%), CD73^+^ (66%). Our data corroborate results published by Pick and colleagues, where cell surface receptors were observed in the CIMVs membranes and their functionality were shown [21]. Kim and colleagues reported that the surface receptors were found similar between MSC-derived microvesicles and MSCs expressing CD13, CD29, CD44, CD73, CD105, CD10 and CD90 [22].

We have demonstrated that CIMVs-MSCs transfer membrane receptors to the target cells. It is known that MSCs surface cell adhesion molecules and signaling receptors play important role in MSCs biology and maintaining the stem like phenotype [27]. The surface receptor transfer by EVs could be the mechanism of mimicry and reprogramming of target cells. Similar data was published by Ratajczak and colleagues, where stem cell-derived microvesicles reprogram target cells by delivering their content including mRNA [26].

MSCs and CIMVs-MSC contained growth factors, cytokines and chemokines, suggesting similar biological activity. To support this assumption, we have demonstrated that human MSCs and CIMVs-MSCs share the angiogenic activity. We have found that human MSCs and CIMVs-MSCs stimulate the sprouting of new blood vessels in vivo. Our data corroborate results published by Gangadaran. et al. where MSCs-derived EVs increased cellular migration, proliferation, endothelial tube formation in vitro and enhanced angiogenesis in ischemic limb *in vivo* [28]. In addition, Lopatina T. et al. reported that MSCs derived EVs could induce the formation of vessel-like structures in vitro and in vivo after the subcutaneous injection in mixture with Matrigel Matrix and human microvascular endothelial cells [29]. We believe that the angiogenic capacity of CIMVs-MSCs depends on growth factors present in their content. Human CIMVs-MSCs demonstrated angiogenic effect in vivo, although it was lower than that of MSCs parental cells. Due to the risks of MSCs therapy connected with undesirable differentiation [4, 5] and transformation [5] the therapeutic use of cell-free therapeutic instrument based on CIMVs is only mechanistically feasible [6]. On the other hand, CIMVs-MSCs could be used to stimulate angiogenesis as they have molecular content and angiogenic activity similar to the parent MSCs. Therefore, CIMVs-MSCs could be used as a method for cell-free regenerative medicine.

## 5. Conclusions

We analyzed the molecular content, receptors expression and angiogenic potential of human MSCs and CIMVs-MSCs. Human CIMVs-MSCs has similar content, immunophenotype and angiogenic activity to that of the parental MSCs. CIMVs-MSCs could transfer membrane receptors to the surface of target cells. Therefore, we believe that human CIMVs-MSCs could be developed for cell-free therapy of degenerative diseases.

## Acknowledgements

We thank the Interdisciplinary Center for Analytical Microscopy, Kazan (Volga Region) Federal University, Kazan, Russia for the conduction of electron microscopy.

## Author Contributions

Study was designed by M.O.G. and A.A.R. M.O.G. conducted characterization of MSCs, CIMVs-MSCs and evaluation of surface receptors transfer. M.N.Z. performed in vivo injections and histology. V.V.V. conducted electron microscopy. I.I.S. and A.V.L. performed proteomic analysis. S.K.K. performed analysis of the subcellular location of proteins using UniProt database. E.V.M. and S.F.K. performed multiplex analysis. L.G.T. isolated MSCs from adipose tissue. J.L.P., A.I.N. did critical revision of the manuscript and N.P.M. edited the manuscript. Data analyses and interpretation were performed by M.O.G. and A.A.R.. Manuscript was written by M.O.G. and reviewed by all co-authors.

## Data Availability

All data generated or analysed during this study are included in this published article (and its Supplementary Information files). The data that support the findings of this study are available from the corresponding author upon request.

## References

1. Rohban, R. and T.R. Pieber, Mesenchymal Stem and Progenitor Cells in Regeneration: Tissue Specificity and Regenerative Potential. Stem Cells Int, 2017. 2017: p. 5173732.

2. Amariglio, N., et al., Donor-derived brain tumor following neural stem cell transplantation in an ataxia telangiectasia patient. PLoS Med, 2009. 6(2): p. e1000029.

3. Rosland, G.V., et al., Long-term cultures of bone marrow-derived human mesenchymal stem cells frequently undergo spontaneous malignant transformation. Cancer Res, 2009. 69(13): p. 5331–9.

4. Kunter, U., et al., Mesenchymal stem cells prevent progressive experimental renal failure but maldifferentiate into glomerular adipocytes. J Am Soc Nephrol, 2007. 18(6): p. 1754–64.

5. Breitbach, M., et al., Potential risks of bone marrow cell transplantation into infarcted hearts. Blood, 2007. 110(4): p. 1362–9.

6. Gomzikova, M.O. and A.A. Rizvanov, Current Trends in Regenerative Medicine: From Cell to Cell-Free Therapy. Bionanoscience, 2017. 7(1): p. 240–245.

7. Yoon, Y.J., O.Y. Kim, and Y.S. Gho, Extracellular vesicles as emerging intercellular communicasomes. Bmb Reports, 2014. 47(10): p. 531–539.

8. Bian, S.Y., et al., Extracellular vesicles derived from human bone marrow mesenchymal stem cells promote angiogenesis in a rat myocardial infarction model. Journal of Molecular Medicine-Jmm, 2014. 92(4): p. 387–397.

9. Haga, H., et al., Extracellular vesicles from bone marrow-derived mesenchymal stem cells protect against murine hepatic ischemia/reperfusion injury. Liver Transplantation, 2017. 23(6): p. 791–803.

10. Deng, M.Y., et al., Mesenchymal Stem Cell-Derived Extracellular Vesicles Ameliorates Hippocampal Synaptic Impairment after Transient Global Ischemia. Frontiers in Cellular Neuroscience, 2017. 11.

11. Peng, L.H., et al., Cell Membrane Capsules for Encapsulation of Chemotherapeutic and Cancer Cell Targeting in Vivo. Acs Applied Materials & Interfaces, 2015. 7(33): p. 18628–18637.

12. Gomzikova, M.O., et al., Cytochalasin B-induced membrane vesicles convey angiogenic activity of parental cells. Oncotarget, 2017. 8(41): p. 70496–70507.

13. Katina M.N., G.R.F., Hayatova Z.G., Emene Ch.Ch., Rizvanov A.A., Isolation, culture and differentiation of rat (Rattus norvegicus) and hamster (Mesocricetus auratus) adipose derived multipotent mesenchymal stromal cells. 2012. VII(3): p. 82–87.

14. Laemmli, U.K., Cleavage of structural proteins during the assembly of the head of bacteriophage T4. Nature, 1970. 227(5259): p. 680–5.

15. Orbay, H., M. Tobita, and H. Mizuno, Mesenchymal stem cells isolated from adipose and other tissues: basic biological properties and clinical applications. Stem Cells Int, 2012. 2012: p. 461718.

16. D. Docheva, F.H.a.M.S., Mesenchymal Stem Cells and Their Cell Surface Receptors Current Rheumatology Reviews, 2008. 4(3): p. 6.

17. Mack, M., et al., Transfer of the chemokine receptor CCR5 between cells by membrane-derived microparticles: a mechanism for cellular human immunodeficiency virus 1 infection. Nat Med, 2000. 6(7): p. 769–75.

18. Flaumenhaft, R., A.T. Mairuhu, and J.E. Italiano, Platelet- and megakaryocyte-derived microparticles. Semin Thromb Hemost, 2010. 36(8): p. 881–7.

19. Todorova, D., et al., Extracellular Vesicles in Angiogenesis. Circ Res, 2017. 120(10): p. 1658–1673.

20. Keshtkar, S., N. Azarpira, and M.H. Ghahremani, Mesenchymal stem cell-derived extracellular vesicles: novel frontiers in regenerative medicine. Stem Cell Res Ther, 2018. 9(1): p. 63.

21. Pick, H., et al., Investigating cellular signaling reactions in single attoliter vesicles. J Am Chem Soc, 2005. 127(9): p. 2908–12.

22. Kim, H.S., et al., Proteomic analysis of microvesicles derived from human mesenchymal stem cells. Journal of proteome research, 2012. 11(2): p. 839–49.

23. Ratajczak, M.Z., et al., Pivotal role of paracrine effects in stem cell therapies in regenerative medicine: can we translate stem cell-secreted paracrine factors and microvesicles into better therapeutic strategies? Leukemia, 2012. 26(6): p. 1166–73.

24. Mussano, F., et al., Cytokine, Chemokine, and Growth Factor Profile Characterization of Undifferentiated and Osteoinduced Human Adipose-Derived Stem Cells. Stem Cells Int, 2017. 2017: p. 6202783.

25. Schinkothe, T., W. Bloch, and A. Schmidt, In vitro secreting profile of human mesenchymal stem cells. Stem Cells Dev, 2008. 17(1): p. 199–206.

26. Ratajczak, J., et al., Embryonic stem cell-derived microvesicles reprogram hematopoietic progenitors: evidence for horizontal transfer of mRNA and protein delivery. Leukemia, 2006. 20(5): p. 847–56.

27. Niehage, C., et al., The cell surface proteome of human mesenchymal stromal cells. PLoS One, 2011. 6(5): p. e20399.

28. Gangadaran, P., et al., Extracellular vesicles from mesenchymal stem cells activates VEGF receptors and accelerates recovery of hindlimb ischemia. Journal of controlled release: official journal of the Controlled Release Society, 2017. 264: p. 112–126.

29. Lopatina T., M.A., Bruno S., Tetta C., Kalinina N., Romagnoli R., Salizzoni M., Porta M., Camussi G., The Angiogenic Potential of Adipose Mesenchymal Stem Cell-derived Extracellular Vesicles is modulated by Basic Fibroblast Growth Factor. Journal of Stem Cell Research & Therapy, 2014. 4(10): p. 1–7.

